# BenchHub enables an inclusive and transparent ecosystem for community-focused benchmarking in computational biology

**DOI:** 10.1101/2025.09.25.678510

**Authors:** Xiaoqi Cabiria Liang, Nick Robertson, Marni Torkel, Sanghyun Kim, Dario Strbenac, Yue Cao, Jean Yee Hwa Yang

## Abstract

The rapid growth of computational methods for the computational biology field highlights the critical role of benchmarking in guiding method selection. However, there is no standardised data structure that effectively links and stores datasets, performance metrics and available ground truth. Without such a unified and shareable structure, it is difficult for the community to contribute, update and extend existing benchmarking studies to ensure long-term relevancy. To address this challenge, we present BenchHub, a community-oriented ecosystem with a modular R6-based structure that enables “living benchmarking.” BenchHub comprises three key components: (i) a Trio database that links datasets, performance metrics, and supporting evidence (e.g. ground truth), (ii) a BenchmarkStudy structure that captures the different benchmark study designs, and (iii) a series of tools together with vignettes and interactive platform that allow users to gain insights from the benchmarking results. Together, these components streamline the benchmarking process for benchmark study developers, methods contributors, and benchmark consumers, promoting reproducibility, comparability, and long-term sustainability in computational biology.

## Introduction

Advances in molecular biotechnology have made it possible to generate vast quantities of high-resolution data (J. Cao et al. 2020),(J. Cao et al. 2020). In response, thousands of analytical methods have been developed to analyse computational biology data and uncover meaningful biological patterns (Zappia, Phipson, and Oshlack 2018). While this growing toolkit provides researchers with analytical flexibility, it also presents a new question: *which method is the most appropriate for which contexts?*

As a result, evaluating method performance has become essential for assessing the quality and reliability of computational tools. This need has driven a surge in benchmarking efforts aimed at comparing methods across diverse tasks and datasets (Y. Cao et al. 2023). In the rapidly evolving field of biology, comparing methods plays a crucial role in helping scientists select the most suitable technique for their research (Dance 2022). Benchmarking not only reveals the strengths and limitations of existing approaches but also helps researchers choose the appropriate tools for their needs.

In response to this demand, the community has introduced sustainable benchmarking practices. Early efforts typically relied on static, one-time comparisons, which posed several challenges. First, these studies were time-intensive to design and replicate (Irizarry, Wu, and Jaffee 2006). Second, while simulated data are widely used to assess method performance, they can introduce systematic bias that may not reflect real-world complexities (Mangul et al. 2019). Third, the choice of evaluation metric can influence the ranking of methods (Norel, Rice, and Stolovitzky 2011). In response, there has been a shift toward creating structured benchmarking frameworks that support continuous updates and community contributions. Unlike one-time comparisons, these frameworks are built around shared infrastructures that integrate standardized datasets, reproducible analysis pipelines, and open contribution mechanisms. Notable examples include Open Problems in Single-Cell Analysis (Luecken et al. 2024), Omnibenchmark (Mallona et al. 2024), and OpenEBench (Capella-Gutierrez et al. 2017), which support ongoing evaluation across diverse research domains.

Despite recent progress, benchmarking studies continue to face several critical challenges. One of the main issues is the lack of ground truth, which makes it difficult to assess method performance (Rawal et al. 2025). In addition, the field lacks a standardised data structure that can effectively link datasets, performance metrics, and ground truth when available. Furthermore, what remains missing is a widely adopted, semi-centralised platform: unlike existing centralised systems that are managed by a single group with full reproducibility, a semi-centralised framework combines the standardization of a centralised system while being flexible by enabling contribution from a wide community.

To address these challenges, we present BenchHub, a semi-centralized framework implemented in R that enables continuous, transparent, and community-driven benchmarking. BenchHub is built around three R6-based components: Trio, BenchmarkStudy and BenchmarkInsights. Together, these components streamline the benchmarking process for three key user groups: developers, contributors, consumers. By promoting standardization, enhancing comparability, and supporting long-term maintenance, BenchHub aims to establish a sustainable infrastructure for benchmarking in computational biology.

## Results

### Results 1 - BenchHub overview, the ecosystem towards living benchmark

BenchHub provides a semi-centralised and community-oriented benchmarking ecosystem that supports “living benchmarking”. It includes three objects: Trio, BenchmarkInsights, and BenchmarkStudy. The Trio object provides a unified data structure that links three key elements in an evaluation or benchmarking study: the datasets used for method evaluation, the metrics for assessing performance, and the supporting evidence, such as ground truth or reference data. Complementing this, BenchmarkInsights enables users to organise benchmarking results and generate informative visualisations to support interpretation and reporting. The third component, BenchmarkStudy, facilitates the design of benchmarking and encourages user contributions (Fig. 1A). Additionally, BenchHub provides a dedicated database to store and share both benchmarking results and the components of Trio objects, including their associated metadata.

**Figure 1.**
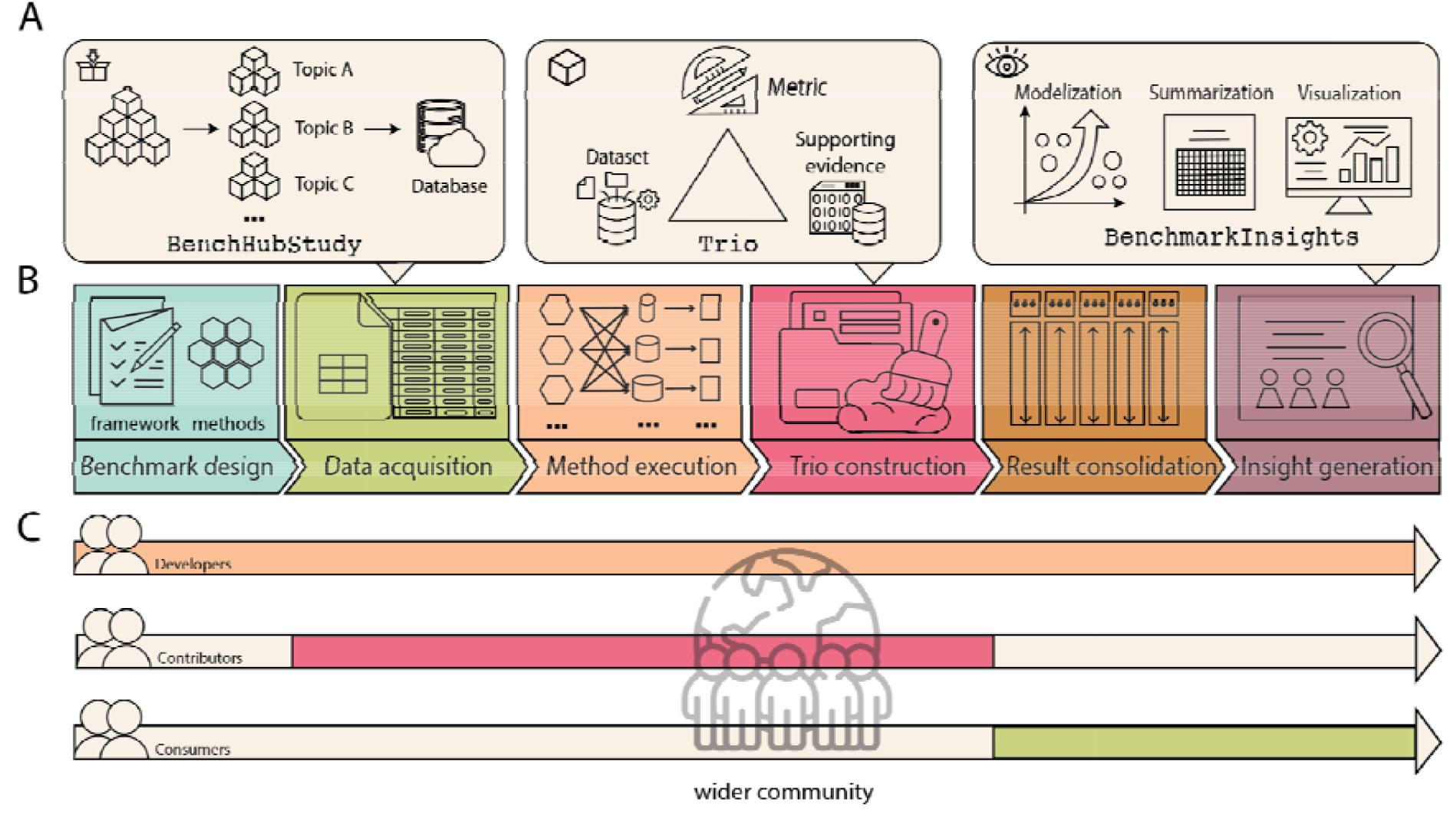
Overview of BenchHub. A. The BenchHub includes three main components. From left to right: BenchmarkStudy, Trio and BenchmarkInsights. B. Benchmark study process includes benchmark design, data acquisition, method execution, object integration, result consolidation, and insight generation. C. Potential users in BenchHub: investigator, method developers and consumers. The grey space represents progress in BenchHub. The three globes in the middle represent work and engagement within a wider community.

BenchHub provides flexibility in how users interact with these three core R objects. While these components are designed to function together, BenchHub’s modular design allows researchers to use its components individually or together, depending on their specific objectives for benchmarking. In other words, different types of “users” of BenchHub will participate at different stages of the workflow (Supplementary Table S1) depending on their requirements. We anticipate at least three main groups of “users”. The first is “benchmark developers”, these researchers will initiate benchmarking studies, typically interact with the full BenchHub workflow, which includes six key steps: (a) study design, (b) data acquisition from BenchHub database, (c) method execution, (d) `Trio` construction, (e) result consolidation, and (f) insight generation (Fig. 1B). They can either start by exploring the BenchHub database and BenchmarkStudy for existing Trio components and framework designs, or build new ones directly. Then they construct custom Trio objects using new datasets, metrics functions and supporting evidence. After generating the benchmark results, they interpret outcomes through the `BenchmarkInsights` object.

The second category is “benchmark contributors,” a broad group of researchers who interact with benchmark studies for various reasons. For instance, method developers often compare their newly developed methods against existing benchmark studies, saving them the effort of conducting a full-scale benchmark themselves. These researchers select the `BenchmarkStudy` object of interest and participate in *method execution* (part c). Here, they apply their method to the collection of `Trio` objects, then *integrate the results* (part d) with the existing data within the BenchmarkStudy object. While their primary goal is likely to demonstrate the superior performance of their methods, their engagement with BenchHub will contribute an additional method to the BenchmarkStudy object. Another example includes researchers who want to re-examine the utility of existing methods for a different purpose. For example, they may explore whether a method designed to analyze gene expression in cancer studies could also be useful for studying another disease, such as diabetes. In this case, they might add new evaluation metrics to assess how well the method works in the new context.

Finally, we have the third category representing “benchmark consumers”, who are researchers interested in the insight generated from the Benchmark studies. Their primary engagement will likely be with part (e) and (f) of the workflow, which mainly involves the `BenchmarkInsights` object. Here, the benchmark consumer will interpret existing results stored in the `BenchmarkInsights` object; perform additional analysis to assess the relative strengths and limitations of available methods; and may add metadata to contextualise datasets or methods without rerunning computations (Fig. 1C). The three R objects and six parts of the workflow represent a modular and flexible design that ensures that BenchHub accommodates a broad range of use cases and levels of technical involvement.

### Results 2 - “Trio”: An R object capturing datasets, evaluation metrics, and supporting evidence information

A robust benchmarking study depends on three components: a dataset for evaluation, clearly defined performance metrics, and supporting evidence such as ground truth or annotations (Fig. 2A). To capture all three components in a given structure, we introduce the concept of a “Trio unit”, which links one dataset (identified by a unique ID), one evaluation metric, and one piece of supporting evidence. Building on this structure, the Trio class is designed to group multiple “Trio unit” objects that share the same dataset ID but differ in evaluation metrics or supporting information. This design enables the user to evaluate a single dataset across diverse benchmarking dimensions, while maintaining modularity and clarity.

**Figure 2.**
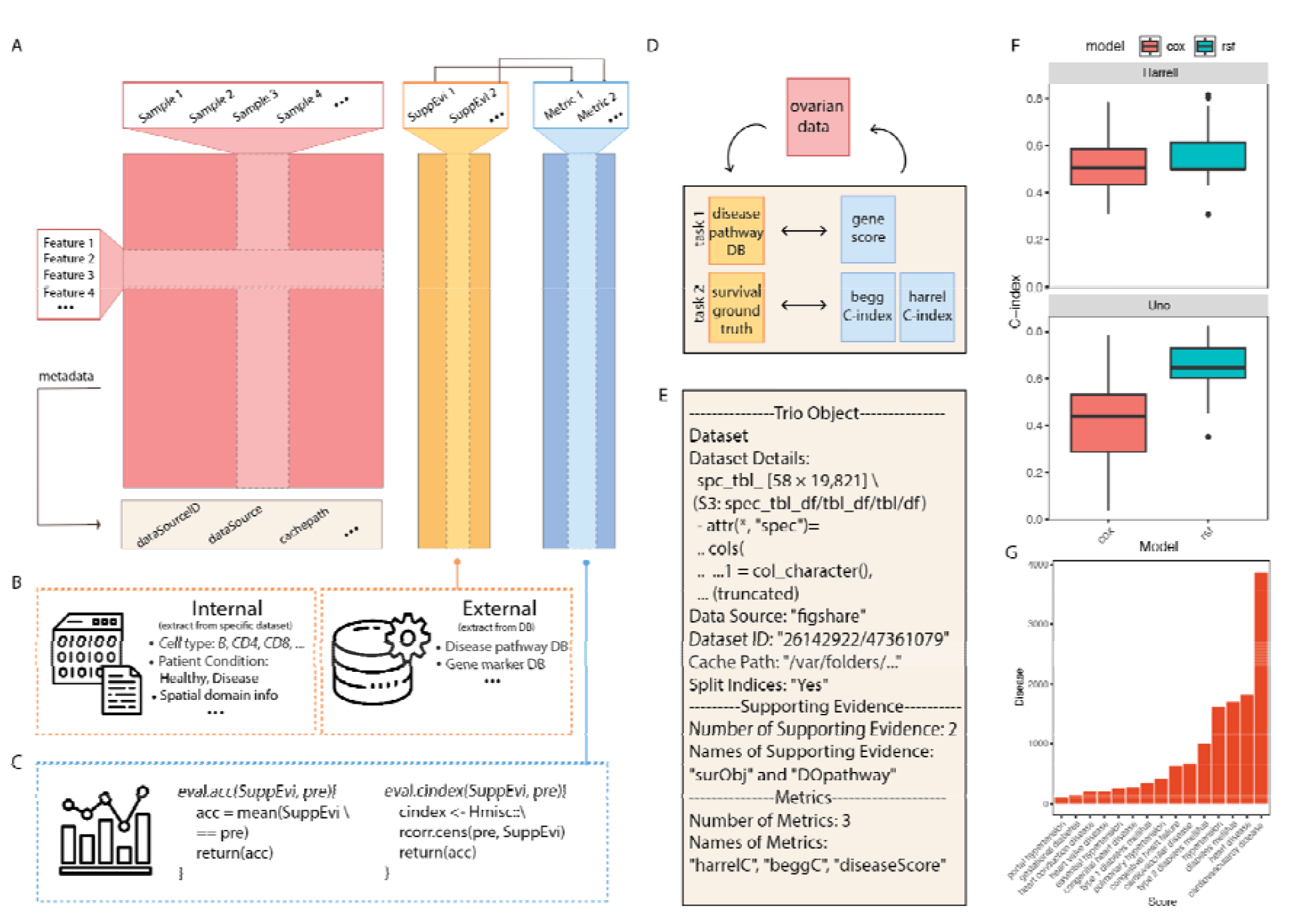
Details of the class structure Trio. **A**. Data structure of Trio includes datasets, supporting evidence, and metrics. Each row of the dataset represents features, and each column represents a sample. Each AuxData and metric are paired combinations, linked together. **B**. The information of supporting evidence can be stored either internally or externally for implementation. **C**. Example pseudocode in metric. The output will be the metric result. **D**. Graphical implementation in survival analysis. **E**. Printing text illustration in Trio. **F**. Boxplots comparing the accuracy of the cox and random forest tree models. The upper panel shows Harrell’s C-index, and the lower panel shows Uno’s C-index. **G**. Top 15 diseases ranked by enrichment score, calculated using gene set enrichment statistics to assess the overlap between dataset-derived gene lists and disease-associated genes from DisGeNET.

Each Trio is composed of three components: the dataset, the evaluation metric, and the supporting evidence. The dataset component refers to the input data used for method evaluation. These datasets are typically hosted in public repositories such as FigShare and can be stored in various file formats, including, but not limited to, CSV, HDF5, or RDS. The second component is supporting evidence, where information can be stored either internally within the Trio object or externally through database links, as illustrated in Figure 2B. Internal supporting evidence commonly includes ground truth labels, such as manual annotations or simulated outcomes. External supporting evidence refers to information retrieved from established databases, which offer biological or clinical context for the dataset under evaluation. The third component, the evaluation metric, defines how performance is measured and compared across methods. These metrics are implemented as user-defined functions, which take as input the supporting evidence and the predicted results, in that order. A simplified pseudocode example of such a function is provided in Figure 2C.

To demonstrate the `Trio` object, we present an example using transcriptomic data from ovarian cancer within a single `Trio` (Fig. 2D). In this case, the dataset serves as the unique identifier, and two pairs of evaluation metrics and supporting evidence are linked through the same `Trio` object (Fig. 2E). The first task involves assessing the predictive performance of two survival models, the Cox proportional hazards model and the survival random forest, using Harrell’s C-index and Uno’s C-index as evaluation metrics, which summarise the agreement between estimated risk scores and all evaluable pairs of patients’ follow-up times. The supporting evidence, representing patient outcomes as a binary event indicator (1 = death or recurrence, 0 = censored), is internally stored as a survival object derived from the original dataset. The resulting C-index scores are summarized in Figure 2F. The second task evaluates disease relevance by applying gene set enrichment statistics to determine how well gene lists derived from the dataset correspond to disease-associated genes. Here, the supporting evidence is retrieved externally from established databases DisGeNET(Piñero et al. 2017), which provides ranked disease gene lists. The top 15 disease scores are shown in Figure 2G. Supplementary Figure S1 presents the pseudocode used to construct this `Trio` object. This example illustrates how a single `Trio` object can be reused across multiple evaluation tasks by linking a dataset to different combinations of metrics and supporting evidence, demonstrating the modularity and extensibility of the `Trio` design.

### Results 3 - Trio ensures consistency in cross-validation evaluations

A key feature of BenchHub is its support for living benchmarks, with results updated as new data or methods become available. A critical aspect of maintaining such a benchmark is ensuring that the cross-validation procedures used to train models and evaluate their accuracy remain comparable. For example, *k*-fold cross-validation is a widely used approach for estimating model performance. It relies on randomly splitting data into training and testing sets. Different repetitions of this process can produce varying data partitions, which may lead to different performance estimates for the same model, if the biological samples are not from a homogeneous population.

We illustrate this with a simulation study in which model performance was assessed using two different data partitions generated by distinct random seeds. Each partition was subjected to five-fold cross-validation, and the average area under the curve (AUC) was calculated for each model. As shown in Figure 3A, substantial differences in performance can emerge between partitions. For example, logistic regression outperformed random forest in the first partition, but the opposite was observed in the second. Comparing the AUC of logistic regression from the first partition with that of random forest from the second could lead to misleading conclusions. In this case, the apparent performance gap reflects variability introduced by data partitioning, rather than true differences between the methods (Fig. 3A).

**Figure 3.**
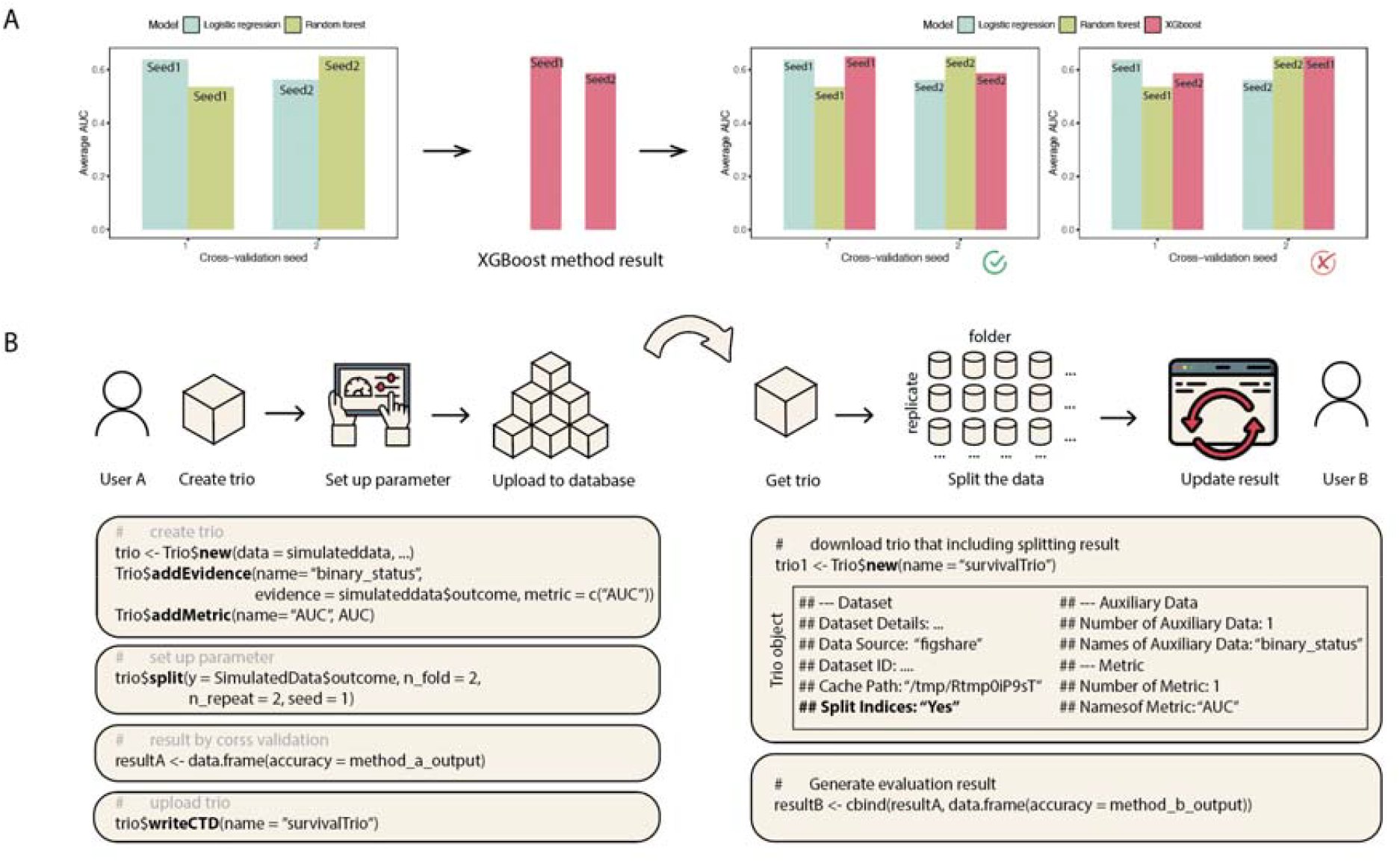
Trio’s ability to handle cross-validation. **A**. Simulation study comparing logistic regression and random forest models across two partition sets. The left one uses the same partition setting but the right one uses the different one. **B**. Detailed schematic workflow. The user A creates a Trio object using supporting evidence and specifies cross-validation parameters (e.g., folds, repetitions, seed). The resulting data splits are saved and uploaded to a shared collection. The user B can then use the same predefined partitions to evaluate new methods, ensuring consistency and comparability across models.

To address this issue, the Trio framework explicitly records and stores the indices of cross-validation splits. As shown in Figure 3B, the process involves two key roles: the original user who creates the Trio object and a subsequent user who wishes to adopt the same cross-validation strategy. The first user begins by contributing a dataset to create a Trio object and specifies parameters such as the number of folds, repetitions, and random seed. The data partition parameter is stored within the Trio object and is uploaded to a shared `BenchmarkStudy` collection. A second user, such as a method developer aiming to compare a new tool with existing results, can retrieve this `Trio` object and access the predefined splitting scheme. This allows them to apply the same training and testing partitions, ensuring consistency in evaluation. For example, using the same simulation study, a user might introduce a support vector machine model. By downloading the original Trio object with its stored parameters, the user can apply the same cross-validation strategy setting. Once the support vector machine is evaluated, its results can be directly compared and combined with those from previously tested models, such as logistic regression and random forest.

### Results 4 - BenchmarkInsights: a structured framework for visualizing and interpreting benchmark results

The third component of BenchHub focuses on extracting insights from benchmarking results (Fig. 4A). To facilitate this, we developed a data structure called BenchmarkInsights, which stores benchmarking outcomes, together with a range of visualisation tools for in-depth analysis. The `BenchmarkInsights` object is initialised using a standardised long-format data frame (Fig. 4B & 4C). The key innovation is its flexibility. These data frames can be generated from evaluation processes within the `Trio` object, or users can supply their own results directly. Additionally, `BenchmarkInsights` supports benchmarking outputs from methods implemented in either R or Python.

**Figure 4.**
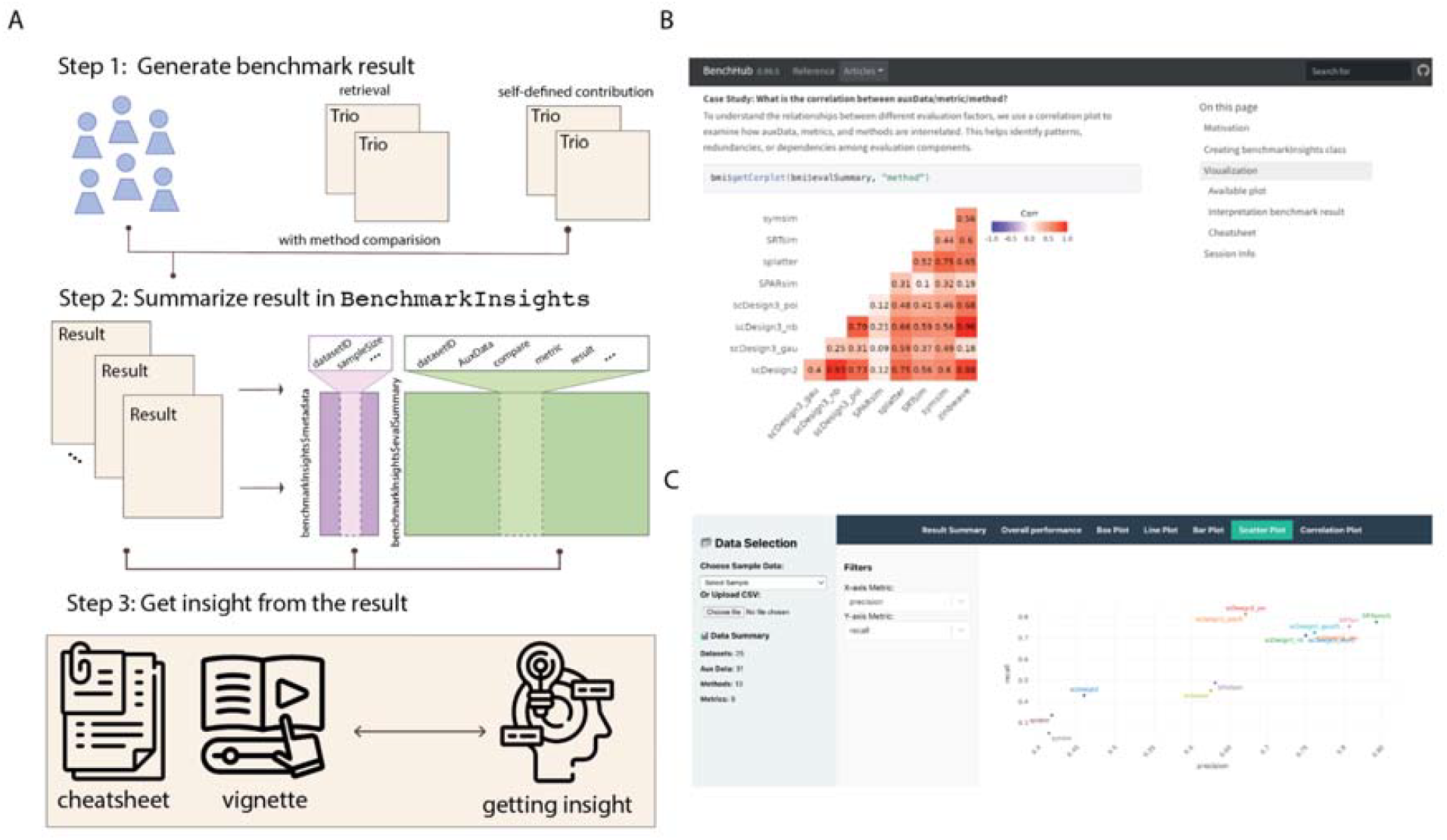
Details in BenchmarkInsights. **A**. Schematic of the data structure of the object BenchmarkInsights. There are two main components: metadata and evalSummary. Metadata includes datasizeID, sampleSize, and additional information. evalSummary is a long-format dataframe. **B**. Vignette examples illustrate a case study, with a correlation plot generated to address the study objective. **C**. An interactive Shiny application with two panels. with the main panel contains seven tabs, each offering different visualisation capabilities.

It is important to note that data structure itself does not generate insights. To bridge this gap, we developed a series of vignettes (https://sydneybiox.github.io/bmi-app/) that serve as a learning resource, demonstrating how users can effectively leverage `BenchmarkInsights` objects to extract relevant information (Fig. 4D). These vignettes are structured around classic benchmarking questions, such as “Which is the best method” (Table 1). Beyond this, we also address more critical and exploratory questions, for example, “Which aspects of the task do all methods fail to capture?” In one vignette, we demonstrate how to approach such questions using linear modelling and visualisation with forest plots. A meta-analysis diagram is used to summarise the relationship between data characteristics and model performance.

**Table 1.** BI case study.

| Case study | Analysis | Function |
| --- | --- | --- |
| What is the overall performance summary? | A heatmap `getHeatmap()` is utilized to provide a high-level overview, summarizing evaluation results across datasets. This visualization quickly highlights general trends and facilitates straightforward comparisons among methods. | `getHeatmap()` |
| What are the correlations between `auxData`, metrics, and methods? | Correlation plots help elucidate relationships among different evaluation factors, revealing patterns, redundancies, or dependencies.<br>To further explore relationships between specific pairs of metrics, scatter plots are employed. These visualizations offer deeper insights into metric alignment or divergence, helping to clarify trade-offs and consistency across methods. | `getCorplot()`<br>`getscatterplot()` |
| What are the computational time and memory trends? | Line plots depict trends in computational efficiency, displaying how methods scale in terms of time and memory usage under varying conditions. These plots highlight potential efficiency trade-offs and identify methods that maintain performance with increasing data complexity. | `getLineplot()` |
| Which metrics most effectively influence | Forest plots quantify and visualize the impact of various metrics on methods, facilitating comparisons that identify the | `getForestplot()` |
| method performance? | most critical evaluation factors. |  |
| How does method variability differ across datasets for specific metrics? | Boxplots illustrate the variability and consistency of methods across multiple datasets for a given metric. This visualisation emphasises robustness or instability, providing valuable insights into method reliability. | `getBoxplot()` |

To enhance accessibility, we have developed an interactive Shiny application, available at (http://shiny.maths.usyd.edu.au/BenchmarkInsights_app/). This platform allows users to upload their benchmarking results and explore insights through an interactive interface. To assist users, a concise cheat sheet is provided to support guide navigation and highlight key features of application (Supplementary Table S2). The interactive platform consists of a menu and result viewer panel (Fig. 4E). In the menu panel, users can either select sample data or upload output from BenchHub to gain insight into the result. The BenchHub output should be CSV format with five required columns: datasetID, method, supporting evidence, metric and result. Once a dataset is selected, a summary appears below, showing the number of datasets, supporting evidence, methods, and metrics included. The result viewer panel contains seven tabs, each providing detailed visualisation capabilities. Additional details are provided in the Methods section.

### Results 5 - Case study: Spatial domain detection

To illustrate the whole BenchHub ecosystem, we present a mini study aimed at benchmarking spatial domain detection. Firstly, the benchmark design step includes two components: selecting methods and defining evaluation strategies. We selected six representative methods: BayesSpace (Zhao et al. 2021), Seurat’s Leiden (Butler et al. 2018), PRECAST (Liu et al. 2023), DR.SC (Liu et al. 2022), BASS (Li and Zhou 2022), and SpatialPCA (Shang and Zhou 2022) (Fig. 5A). The evaluation strategy was structured around three categories of criteria: accuracy, spatial continuity, and scalability, with each category assessed using multiple metrics (see Material & Methods).

**Figure 5.**
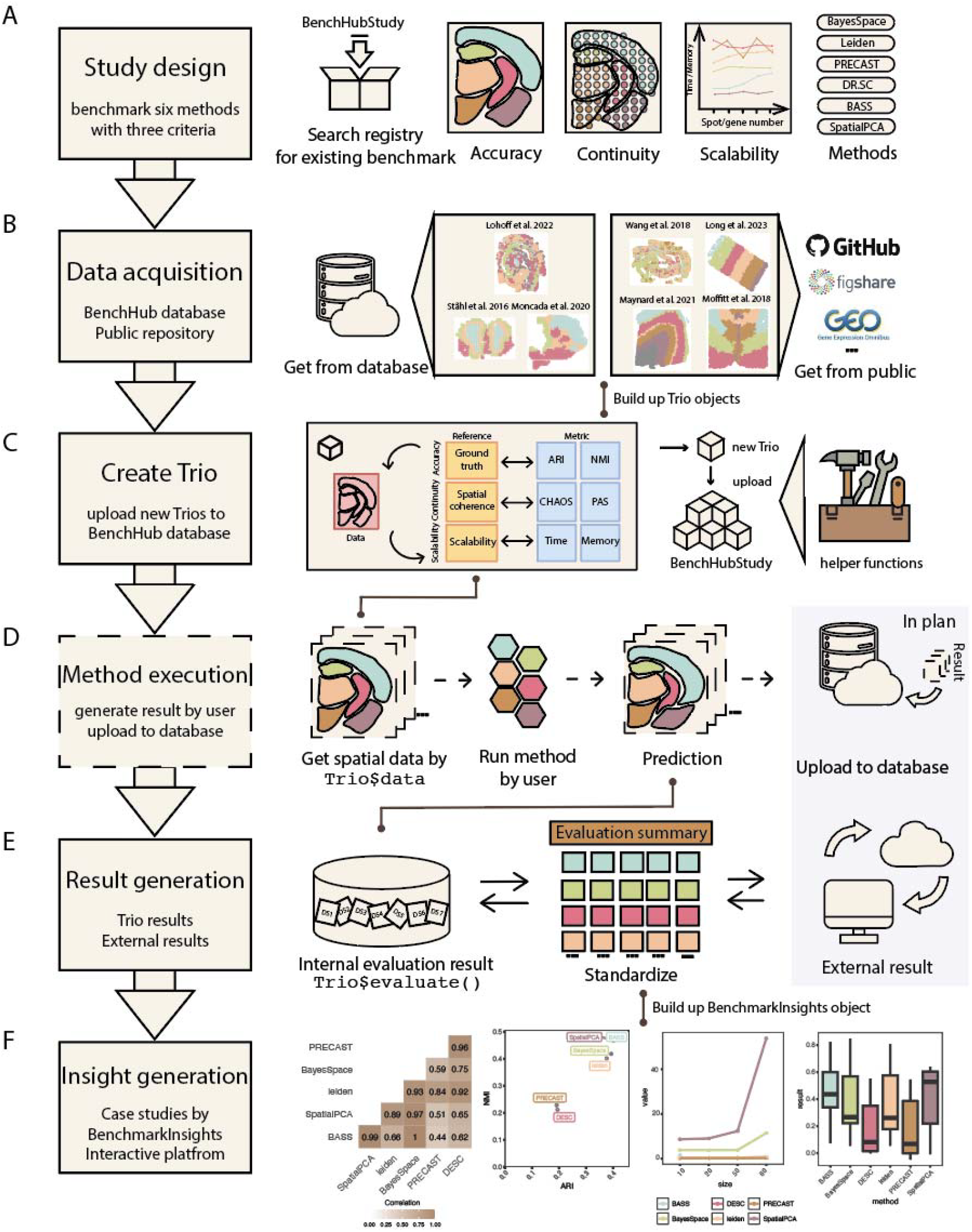
Spatial domain detection case study. A. Benchmark study design. It includes the evaluation tasks and spatial domain detection methods. B. Datasets used in this benchmark study. Difference colours represent different spatial clustering. C. Method execution. We systematically applied the selected spatial domain detection methods across the acquired datasets to generate predicted spatial domains. D. Trio integration. The basic structure of the trio used in this study. E. Result consolidation. The evaluation result for each dataset is stored in long format. F. Insight generation. Using the BenchmarkInsights object, we can get insight from the evaluation result automatically. G. User contribution. All the trios would upload to the BenchmarkStudy collections including the helper function used in this case study.

Based on our study design, we first searched the BenchmarkStudy registry and confirmed that no existing entries aligned with our benchmarking objectives. We then explored the BenchHub database and identified three datasets with manual annotations that were suitable for our analysis. To complement these, we collected additional four publicly available spatial transcriptomics datasets across diverse sequencing protocols and tissue types from both human and mouse sources (Fig. 5B). Each dataset contains manual annotations, which are used as ground truth for benchmarking. For each dataset, we constructed a corresponding Trio object, resulting in a total of seven Trio objects, each uniquely identified by its dataset ID (Fig. 5D). Each Trio stored three pairs of supporting evidence and evaluation metric functions. The first pair linked manual annotations with average rate index (ARI) and normalized mutual information (NMI) to assess clustering accuracy. The second captured spatial coherence through spatial chaos score (CHAOS) and proportion of abnormal spots (PAS) scores, calculated directly from the dataset. The third pair evaluated scalability using runtime and memory consumption metrics. To generate spatial domain predictions, we applied a set of selected spatial domain detection methods (Fig. 5C).

During result consolidation, predicted spatial domains were compared with manual annotations to compute evaluation metrics using the predefined functions. Scalability results were manually integrated into the corresponding Trio objects. All benchmarking results were stored in a long-format structure (Fig. 5E). Using the consolidated benchmark results, we initialized benchmarkInsight objects for interpretation. We examined several analyses, including: a summary overview; correlations among supporting evidence, metrics, and methods; trends in computational efficiency; identification of the most effective metric for each method; and examination of method performance variability across different datasets for specific metrics (Fig. 5F).

Here writing upload into the bench hub database or like BenchmarkStudy object, pending To support user interaction and contribution within the benchmarking ecosystem, we used a BenchmarkStudy object in combination with the BenchHub database to organize all Trio objects associated with a given benchmarking task. For example, in the spatial domain detection study, seven Trio objects were created, each corresponding to a different dataset. We uploaded the associated datasets, evaluation metrics, supporting evidence, and a set of helper functions designed to facilitate evaluation (Fig. 5G & SuppFig. S2). One such helper function filters out low-quality spatial spots prior to applying domain detection methods. The BenchmarkStudy object links all these components under the relevant task category, in this case spatial domain detection.This structure allows benchmark contributors to retrieve existing BenchmarkStudy objects, evaluate new methods using consistent inputs, and contribute additional results.

## Discussion

Here, we present a benchmarking framework (BenchHub), built on the Bioconductor ecosystem, to support the evaluation of novel computational methods in the computational biology field. The motivation arises from an emerging data science challenge, where the exponential growth of computational tools addressing similar tasks necessitates a quantitative decision framework to identify and select the most appropriate method for a given context. Our proposed solution adopts a semi-centralized system with a modular R6-based structure and serves as a way to implement the concept of living benchmarking. The framework comprises three distinct components: a Trio data structure to manage evaluation datasets and metrics, a BenchmarkStudy module for method evaluation, and a benchmarking insight layer for analyzing the results.

The primary advantage of R6 is that it follows the encapsulated object-oriented programming (EOOP) paradigm. EOOP incorporates several features that have been used for the creation of BenchHub. R6 implements EOOP modularity, meaning that an R6 object is totally self-contained. Everything about its state, all its methods, and its data are fields of the object itself. This enables the crucial implementation of the BenchHub class, where each benchmark study, such as the spatial simulation study, is represented as a distinct object. All the relevant components, such as study description, the list of trios and evaluation functions, are stored within this object. This self-contained design ensures that as the number of benchmark studies in BenchHub grows, they can be stored in an organised and scalable fashion. Lastly, EOOP is the dominant paradigm outside R such as Python and C++, meaning that we can target a broad audience of people that are not necessarily familiar with R’s many different and idiosyncratic OOP systems such as S3 or S4.

The concept of a “living benchmark” is gaining traction in computational biology and bioinformatics, with several notable implementations. Open Problems in Single Cell Analysis uses a centralised platform where datasets, methods, and tasks are evaluated via a standardised pipeline. Similarly, Polaris (for drug discovery) offers centralised access to data and evaluation tools through an API. Our work, BenchHub, extends a semi-centralised approach that was pioneered by Affycomp (Irizarry, Wu, and Jaffee 2006) where users download standard data, run methods locally, and submit results for comparison. BenchHub extends such an approach by creating a repository for collecting evaluation data, enabling contributors to run workflows locally and share results within a modular benchmarking framework. This semi-centralised design allows greater flexibility as it allows users to contribute locally using their own computational environment without the additional learning hurdle, while ensuring the data and metadata are centralised and standardised. Through this flexibility, BenchHub aims to lower barriers to user contribution and encourage greater community engagement.

BenchHub’s flexibility allows it to complement and integrate with other existing strategies. In particular, when working with centralised systems like OpenProblems, BenchHub can engage in two distinct ways. First, the BenchHub Trio design includes a collection of evaluation datasets with corresponding supporting evidence, such as, but not limited to ground truth labels. These curated datasets can be incorporated into the OpenProblems system to their existing benchmarking tasks and add to their collection of evaluation results. Second, the results from these evaluations can be imported back into BenchHub’s BenchmarkInsight component for more flexible, downstream analysis if needed. Third, BenchmarkInsight takes any evaluation results into a plain data frame format. Any evaluations from the centralised systems such as OpenProblems can be imported into the BenchmarkInsights for further in-depth exploration of results.

While benchmarking provides a structured framework to compare computational methods, it does not replace critical thinking that is required for the effective design of benchmarking studies. Designing meaningful benchmark studies involves careful consideration of what is being measured and its context. For example, accuracy may be insufficient in imbalanced datasets, where metrics like sensitivity or specificity might be more informative. The field of single-cell research recently questioned the use of overall correlation between predicted and measured gene expression intensities as an evaluation metric (C. Wang et al. 2025). They noted that when the gene expression vector includes many non-expressed genes, including comparison of white noise in the average correlation can be misleading. Instead, they proposed computing correlations using only highly variable genes (or “signal genes”) to better reflect biological relevance and prediction performance. BenchHub supports benchmarking, but the design of each study remains an important scientific component to BenchHub.

The establishment of BenchHub generates the information needed to answer our initial question: which method is most appropriate for which context? Going forward, this paves the way for a more automated, data-driven method recommendation system, where we need to develop a strategy to (i) identify or match relevant data characteristics, and (ii) select and define relative weights of a set of evaluation criteria to inform overall method ranking. The success of BenchHub will rely on community adoption and, in particular, the quality of contributions. As new studies are contributed by users, it is important to maintain consistency and high standards of the contributions. To achieve this, we plan to engage the community through hands-on training at national and international conferences.

In summary, BenchHub, a semi-decentralsed framework with structured foundation (Trio) and modular organisation system (BenchmarkStudy object) and benchmark result interpretation (BenchmarkInsights object), this implementation reinforces the principle of *living benchmarking* by enabling continual updates and analysis across evolving datasets and methods. It (i) facilitates access to ready-made datasets and ground truth resources for benchmarking, and (ii) enables straightforward evaluation of new tools by method developers. Together, these components support the storage, sharing, and access of benchmarking resources in a consistent and interoperable format.

## Material & Methods

### Trio object

The concept of “Trio” is built around three key components: a dataset, a metric, and supporting evidence. The basic unit within this framework, called a “Trio unit”, is implemented as an R data object. Each “Trio unit” contains one dataset, one metric, and one supporting evidence. A trio object can contain multiple metrics and supporting evidence for one dataset. When inspecting a trio object, it displays three main sections:

- Dataset: includes details such as the data source, dataset ID, cache path, and split indices.s
- Metrics: shows the number and names of metrics defined within the object.
- Supporting evidence: indicates the number and names of supporting evidence included.

A trio object can be instantiated directly using the `Trio`$`new()` constructor through three main approaches:

- Curated trio datasets: utilizing predefined datasets listed in the curated trio datasets collection. This method allows rapid initialization with pre-populated metrics and supporting evidence, suitable for quick evaluations.
- Source and ID: creating an object by specifying a dataset ID from an external source that supports a valid trio downloader (e.g., Figshare, GEO, or ExperimentHub). This method is particularly useful when working with known external dataset identifiers.
- Direct object loading: passing a dataset already loaded into the R environment directly into the constructor. This approach is convenient when working with locally prepared datasets.

Evaluation metrics are integrated into the Trio as pairwise functions of the form `f(expected, predicted)`, returning a single value. Additional parameters can be provided through the `args` argument as a named list. Supporting evidence can be added either as fixed values or as functions that dynamically compute values from the dataset.

Once relevant metrics and supporting evidence are defined, evaluations can be executed using the `evaluate()` function. This function supports both simple and multi-method evaluations. To improve efficiency, Trio incorporates caching to eliminate repeated lengthy downloads. Data accessed initially is stored locally, with the cache directory path specified via the `cachePath` parameter. If unspecified, it defaults to `∼/.cache/R/TrioR/`.

Trio also supports robust data splitting for cross-validation purposes. The split method divides datasets into training and test subsets, with indices for each split stored in the `splitIndices()` attribute. These indices are generated using the `splitTools` package, allowing specification of the outcome variable, the number of folds, and repetitions.

Furthermore, the outcome variable can be stratified using the `stratify` parameter to ensure balanced splits.

### BenchmarkInsights object

The BenchmarkInsights (BI) container class is another R6-based R data object within BenchHub to support analysis and visualization of benchmark results. BI operates on a standardized input data frame, typically generated from Trio evaluations. BI implemented a range of functions that allow users to interact with and explore benchmarking outcomes from multiple perspectives, such as comparison of methods, relationship between evaluation metrics and factors affecting method performance BI supports the update of evaluation results through the `addEvalSummary()`, which enables users to append new evaluation results onto existing one. The `addMetadata()` function allows the inclusion of method-specific metadata, to support context-specific interpretation of results.

BI includes six visualization functions, five illustrative case studies, a user cheat sheet, and an interactive ReactJS application. Key visualisations include heatmap, correlation plot, boxplot, forest plot, scatter plot and line plot. Detailed descriptions of the available plots and case studies are provided in Table 1. The BI cheat sheet is available in Supplementary Table S2. The ReactJS app, designed to support exploratory analysis of BI objects, is accessible online at: https://sydneybiox.github.io/BI-app/.

### BenchmarkStudy object

Benchmarking studies requires multiple Trios. To facilitate the organisation of Trios in benchmarking studies, we designed a BenchmarkStudy object. The BenchmarkStudy object contains the following core components: i) collection of Trio IDs used in benchmarking studies, ii) mapping functions that format the method output for evaluation functions and iii) metadata containing description of the benchmarking study.

In terms of storing Trios, BenchmarkStudy stores the ID rather than the Trios. When a user downloads a study, the relevant Trios are then downloaded in real time by referencing the database. This design is to ensure scalability of the storage, especially when benchmarking studies can sometimes involve over 100 datasets (i.e., Trios) and that each Trio could be large. This also enables convenient updating of a benchmarking study. For example, when new Trios get included, a new version can be created with the updated Trio ID, without needing to upload another bulky BenchmarkStudy object and causing storage strain.

Mapping functions are post-processing helper functions that format method outputs into a standard format for evaluations. The purpose of mapping functions is to store the script that a benchmarker used and to allow reproducibility in evaluation. Using simulation method benchmarking as an example. An example of a mapping function is a function that computes the sparsity of the data. These mapping functions can be run on data generated by a simulation method and real data to compare whether the sparsity of the simulated data mimics that of the real data.

### BenchHub Database

To support community-driven benchmarking, BenchHub organises its data structure using the Curated Trio Datasets Google Sheet. It comprises five sheets: Datasets, Dataset-AuxData, TaskAuxData, Task Metric, Metric, and AuxData Type. Users can contribute to or modify entries using the `writeCTD()` function, which simplifies the submission process. Before using this function, contributors must generate a personal access token on GitHub and authenticate access to the shared Google Sheet. Alternatively, users may manually edit or append new entries directly within the spreadsheet.

The contribution process begins by uploading the dataset associated with a Trio to a supported public repository such as Figshare, Zenodo, or GEO. Once the dataset is hosted online, users register the corresponding Trio entry in the BenchHub database using the `writeCTD()` function. This function collects essential metadata, including the dataset name, a direct download link, and whether the dataset is omics-based or clinical. It then automatically opens a GitHub page on the user’s local machine, where evaluation metric functions can be uploaded. By managing data and metrics as separate components, BenchHub offers flexibility in contribution: users with datasets can register them independently, while others can provide evaluation metrics through GitHub without needing access to the raw data. When a user calls `Trio`$`new()`, the object is dynamically constructed by retrieving the dataset from the provided URL and downloading the corresponding evaluation metrics from the GitHub repository.

### Case study I: Simulation study

#### Dataset description

To simulate a classification task with class imbalance, we generated a synthetic dataset with 200 samples and 20 features. The feature matrix was sampled from a multivariate normal distribution with mean zero and identity covariance. The binary outcome variable was created with a 10% positive class proportion, introducing a controlled imbalance representative of many real-world biomedical datasets.

#### Benchmark method

We compared two commonly used classification algorithms: logistic regression and random forest. Logistic regression was implemented using a generalized linear model with a binomial link function. The random forest classifier was trained using default parameters. We use two different seeds to perform the training and testing splitting. For each setting, the seed was used to generate a distinct set of five fold cross validation partitions. The average AUC across the five folds was computed for each model. Presenting results from two different seeds illustrates how variation in the data partitioning can influence model performance and ranking.

#### Evaluation of accuracy

Model performance was assessed using the area under the receiver operating characteristic curve (AUC). For each data partition, five-fold cross-validation was applied, and the mean AUC across the five folds was recorded.

- AUC: measures a model’s ability to discriminate between binary outcomes.

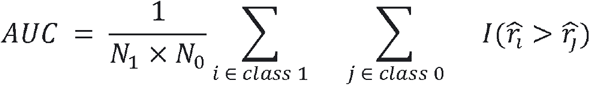

Where *N*_1_ and *N*_0_ are the number of positive and negative instances. 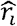 and 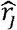 are the predicted probability for a positive sample *i* and a negative sample j. *I*(.) is the indicator function that returns 1 if the condition holds and 0 otherwise.

### Case study II: Survival analysis

#### Dataset description

For this study, we used a publicly available ovarian cancer transcriptome dataset referred to as ovarian in this paper (Ganzfried et al. 2013). The dataset includes 58 observations (patient samples) and 19,818 genes. The event of interest is subject to censoring, with a censoring rate of 38%.

#### Benchmark method

We benchmarked two survival analysis methods: the cox proportional hazards model and the random survival forest. The cox model is a semi-parametric approach that estimates the hazard ratio based on covariates under the proportional hazards assumption. In contrast, the random survival forest is a non-parametric ensemble learning method that can capture interactions and nonlinear effects without relying on proportional hazards.

#### Evaluation of accuracy

To evaluate the accuracy of survival models, we apply two metrics: Harrell’s C-index and Uno’s C-index. These metrics assess the models’ ability to rank survival times based on predicted risk scores.

- Harrell’s C-index: measures the concordance between predicted risk scores and observed survival times.

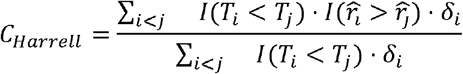

Where T_*i*_ is the observed time, *δ*_*i*_ is the event indicator *I*(.) (1 if event occurred, 0 if censored), and 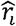 is the predicted risk score for subject *i*
- Uno’s C-index: measures the discriminative ability of a survival model while accounting for censoring bias.

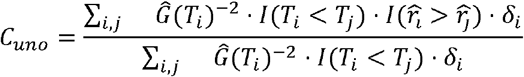

Where 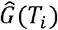 is the kaplan-meier estimate of the censoring distribution at time *T*_*i*_,*δ*_*i*_ is the event indicator *I*(.) (1 if event occurred, 0 if censored), and 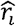 is the predicted risk score for subject

#### Evaluation of gene score

We evaluated disease gene scores using gene set enrichment statistics to quantify the relevance of selected gene lists to known disease-associated genes. As an external reference, we used DisGeNET(Piñero et al. 2017), a curated database of disease–gene associations. Gene scores were compared against disease gene rankings from DisGeNET(Piñero et al. 2017), and the top 15 enriched disease gene sets were summarized.

### Case study III: Spatial domain detection

#### Dataset description

For this benchmark study, we compiled seven spatial transcriptomics datasets, each manually annotated with spatial domain labels. The collection spans a range of experimental protocols and includes tissue samples from both human and mouse sources. Full details for each dataset are provided in Supplementary Table S3.

#### Benchmark method

We selected six widely recognized spatial domain detection methods based on literature review: BayesSpace (Zhao et al. 2021), Seurat’s Leiden (Butler et al. 2018), PRECAST (Liu et al. 2023), DR.SC (Liu et al. 2022), BASS (Li and Zhou 2022), and SpatialPCA (Shang and Zhou 2022). Detailed information on these methods, including software versions, associated references, and default parameter settings, is provided in Supplementary Table S4.

#### Evaluation of accuracy

Spatial domain detection involves grouping spatial spots based on similar expression patterns. To evaluate the accuracy of spatial clustering results, we apply two metrics: Adjusted Rand Index (ARI) and Normalized Mutual Information (NMI). These metrics assess the agreement between manually annotated ground truth clusters and predicted spatial domain clusters.

- Adjusted rand index (ARI): measures the similarity between the ground truth and predicted clustering outcomes. The ARI is defined as:

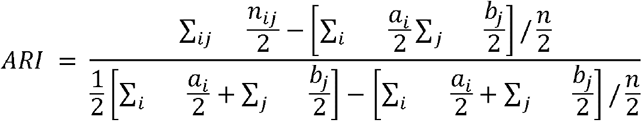

where *n* is the total number of spots; *a*_*i*_ is the number of spots in the *i*^*th*^ cluster of the ground truth; *b*_*j*_ is the number of spots in the *j*^*th*^ cluster of the predicted clustering; *n*_*ij*_ is the number of spots that are in both the *i*^*th*^ ground truth cluster and the *j*^*th*^ predicted cluster.
- Normalized mutual information (NMI): quantifies the mutual dependence or shared information between the ground truth and predicted clustering results. NMI is defined as:

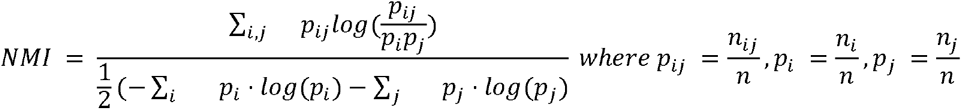

where *n* is the total number of spots; a_*i*_ is the number of spots in the *i*^*th*^ cluster of the ground truth; *b*_*j*_ is the number of spots in the *j*^*th*^ cluster of the predicted clustering; *n*_*ij*_ is the number of spots that are in both the *i*^*th*^ ground truth cluster and the *j*^*th*^ predicted cluster.

Evaluation of continuity

To evaluate the continuity of spatial clustering results, we apply two metrics: Proportion of Abnormal Spots (PAS) and spatial chaos score (CHAOS).

- Spatial chaos score (CHAOS): measures the spatial continuity of the detected spatial domains (Alexandrov and Bartels 2013) (Guo et al. 2021). To compute CHAOS, we first construct a one-nearest-neighbor (1NN) graph for spots within each spatial cluster, linking every spot to its closest neighbor. CHAOS is defined as:

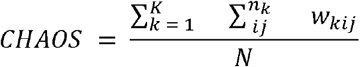

where *n*_*k*_ is the number of cells in the *k*^*th*^ spatial domain; *n* is the total number of cells in the data; and *K* is the number of unique spatial domains; *w*_*kij =*_ *d*_*ij*_ if *spot*_*i*_ and *spot*_*j*_ are connected in the 1NN graph in cluster *k* and 0 otherwise.
- Proportion of Abnormal Spots (PAS) (Shang and Zhou 2022): evaluates the randomness or inconsistency of spot assignments relative to their surrounding neighborhood. It calculates the proportion of spots whose assigned cluster labels differ from at least six out of their nearest neighbors. A low PAS score indicates high homogeneity within spatial clusters. PAS is defined as:

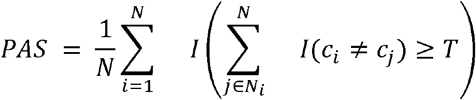

where N is the total number of spots; N_*i*_ denotes the set of nearest neighbors of spot *i*;*c*_*i*_ is the cluster label of spot *i*, and *c*_*i*_ is the cluster label of neighbor *j*; *T* is the threshold number of mismatched neighbors and *I*(.) is the indicator function, returning 1 is the condition is true, and 0 otherwise.

#### Evaluation of scalability

Our scalability analysis was limited to Dataset 6, which we systematically downsampled to generate datasets with different numbers of spots and genes. Specifically, we considered spot counts of 1,000 and 2,000, and gene counts of 1,000 and 2,000, resulting in four distinct downsampled datasets.

The execution time for each method was measured using the bench package in R, specifically using the `bench::mark()` function with memory profiling enabled. Each method was executed three times to ensure reliability, and the median execution time and allocated memory were calculated. Methods encountering errors during execution were captured and logged accordingly, with unsuccessful runs excluded from final summarization. Memory consumption was expressed in megabytes, and execution times were averaged across repetitions to ensure accuracy.

#### Computational resources

Computational benchmarking on an R server equipped with Intel Core i9-14900K CPUs (5.2 GHz, 36 MB Smart Cache, 24 cores in total) and 64 GB of DDR5 memory operating at 6000 MHz.

## Data Availability

All datasets used in this study are publicly available and were downloaded from the following links: (1) Dataset 1 reported in (Ståhl et al. 2016). Mouse olfactory bulb. Downloaded from the supplementary section of the corresponding paper (www.spatialtranscriptomicsresearch.org); (2) Dataset 2 reported in (Moncada et al. 2020). Human pancreatic ductal adenocarcinomas. Downloaded from GEO accession GSE111672 https://www.ncbi.nlm.nih.gov/geo/query/acc.cgi?acc=GSE111672; (3) Dataset 3 reported in (Lohoff et al. 2022). Mouse embryo. Downloaded from https://marionilab.cruk.cam.ac.uk/SpatialMouseAtlas/; (4) Dataset 4 reported in (X. Wang et al. 2018). Mouse medial prefrontal cortex. Downloaded from http://clarityresourcecenter.org/; (5) Dataset 5 reported in (Long, Miller, and The SpaceTx Consortium 2023). Mouse primary cortex. Downloaded from https://spacetx.github.io/; (6) Dataset 6 reported in (Maynard et al. 2021). Human postmortem DLPFC. Downloaded from http://research.libd.org/spatialLIBD; (7) Dataset 7 reported in (Moffitt et al. 2018). Mouse preoptic hypothalamus. Downloaded from https://doi.org/10.5061/dryad.8t8s248

## Code Availability

The BenchHub software code is publicly available at https://sydneybiox.github.io/BenchHub/. The source code is released at https://github.com/SydneyBioX/BenchHub/. Example codes for using BenchHub are publicly available at https://sydneybiox.github.io/BenchHub/index.html. The BenchHub also provides an interactive platform for users to upload and explore benchmark evaluation results. This web tool, developed using ReactJS is available at https://sydneybiox.github.io/bmi-app/.

## Acknowledgments

The authors thank all their colleagues, particularly at The University of Sydney, Sydney Precision Data Science and Charles Perkins Centre for their support and intellectual engagement. Special thanks to Ellis Patrick, Shila Ghazanfar and Chunhan Wang for their contributions in discussions.

## Author information

### Author contributions

J.Y.H.Y. and Y.C. conceived and led the study. X.L. ran the case studies and performed interpretation of the evaluation framework with close guidance from J.Y.H.Y. and Y.C.. M.T. designed BenchmarkInsights web tool. The implementation and construction of the BenchHub R package for the case study were done jointly by N.R., X.L., Y.C. D.S. and S.K.. The development of the designed template was done jointly by all authors and all authors wrote, reviewed and approved the manuscript.

### Authors and Affiliations

School of Mathematics and Statistics, The University of Sydney, NSW 2006, Australia. Xiaoqi Liang, Nick Robertson, Marni Torkel, Sanghyun Kim, Dario Strbenac, Yue Cao & Jean Yee Hwa Yang

Sydney Precision Data Science Centre, The University of Sydney, NSW 2006, Australia. Xiaoqi Liang, Nick Robertson, Marni Torkel, Sanghyun Kim, Dario Strbenac, Yue Cao & Jean Yee Hwa Yang

Charles Perkins Centre, The University of Sydney, NSW 2006, Australia. Xiaoqi Liang, Nick Robertson, Sanghyun Kim, Dario Strbenac, Yue Cao & Jean Yee Hwa Yang

Business School, The University of Sydney, NSW 2006, Australia. Sanghyun Kim

## Ethics declarations

### Ethics approval and consent to participate

Not applicable

### Consent for publication

Not applicable

