## Supplementary materials for "BenchHub enables an inclusive and transparent ecosystem for community-focused benchmarking in computational biology"

**Supp Table S1.** Details of the six elements workflow

| Workflow elements | Details |
| --- | --- |
| Study design | Users initiate the process by defining the benchmarking framework and selecting a series of methods relevant to their analysis goals. Here we select the (i) Methods we like to compare (ii) the metrics we intend to use and (iii) consider the types of datasets we need. |
| Data acquisition | A number of data acquisitions (i) Selecting from the database, users will search the BenchmarkStudy registry for existing benchmarking entries that align with their study and download relevant data (Trio) or datasets. (ii) Identify datasets from public repositories or their own experimental data; (iii) Create simulated data. |
| Trio construction | If relevant Trios are available, they can be downloaded from the BenchHub database. Otherwise, users may construct their own Trio objects by linking datasets, methods, and ground truth or supporting evidence. |
| Method execution | With both datasets and methods in place, users independently execute the selected methods to generate predictions or analysis results. |
| Benchmark result generation | Results refers to “evaluation results” not output from methodology. The evaluation() function can be used on a Trio object to compute benchmarking results. This function returns a standardised long-format dataframe containing evaluation metrics. |
| Insight generation | Benchmarking results can be used to create a benchmarkInsight object, which enables users to explore, interpret, and visualise performance trends and insights across datasets and methods. |

**Supp Table S2.** Details of the cheatsheet.

| **Question** | **Code** |
| --- | --- |
| Summary overview | getHeatmap(evalReuslt) |
| Correlation analysis | getCorplot(evalReuslt, input_type) |
| Scalability trend (time/ memory) | getLineplot(evalReuslt, order) |
| Metric-Model impact (Modeling) | getForestplot(evalReuslt, input_group, input_model) |
| Method variability across datasets | getBoxplot(evalReuslt) |
| Metric relationship | getScatterplot(evalReuslt, variables) |

**Supp Table S3.** Details of the spatial data in the case study.

| **Dataset** | **Species** | **Tissue** | **Protocol** | **Spot number** | **Gene number** | **Ref** | **Download** |
| --- | --- | --- | --- | --- | --- | --- | --- |
| **Dataset 1** | Mouse | Olfactory bulb | ST | 278 | 182 | Ståhl et al. 2016 | [link](http://www.spatialtranscriptomicsresearch.org) |
| **Dataset 2** | Human | Pancreatic ductal adenocarcinomas | ST | 428 | 25753 | Moncada et al. 2020 | [link](https://www.ncbi.nlm.nih.gov/geo/query/acc.cgi?acc=GSE111672) |
| **Dataset 3** | Mouse | embryo | seqFISH | 8425 | 351 | Lohoff et al. 2022 | [link](https://marionilab.cruk.cam.ac.uk/SpatialMouseAtlas/) |
| **Dataset 4** | Mouse | Medial prefrontal cortex | STARmap | 1049 | 166 | Wang et al. 2018 | [link](http://clarityresourcecenter.org/) |
| **Dataset 5** | Mouse | Primary cortex | BaristaSeq | 2042 | 79 | Long et al. 2023 | [link](https://spacetx.github.io/) |
| **Dataset 6** | Human | Postmortem DLPFC | 10x Visium | 33538 | 3639 | Maynard et al. 2021 | [link](http://research.libd.org/spatialLIBD) |
| **Dataset 7** | Mouse | Preoptic hypothalamus | MERFISH | 5488 | 155 | Moffitt et al. 2018 | [link](https://doi.org/10.5061/dryad.8t8s248) |

**Table S4.** Details of the spatial clustering methods evaluated in this study.

| **Methods** | **Year of publication** | **Package version** | **Parameter setting** |
| --- | --- | --- | --- |
| **BayesSpace** | 2021 | 1.14.0 | The number of spatial domains sets to the true number; others are recommended settings. |
| **Leiden by Seurat** | 2018 | 5.1.0 |  |
| **PRECAST** | 2023 | 1.6.5 |  |
| **DR.SC** | 2022 | 3.4 |  |
| **BASS** | 2022 | 1.3.1 |  |
| **SpatialPCA** | 2022 | 1.3.0 |  |

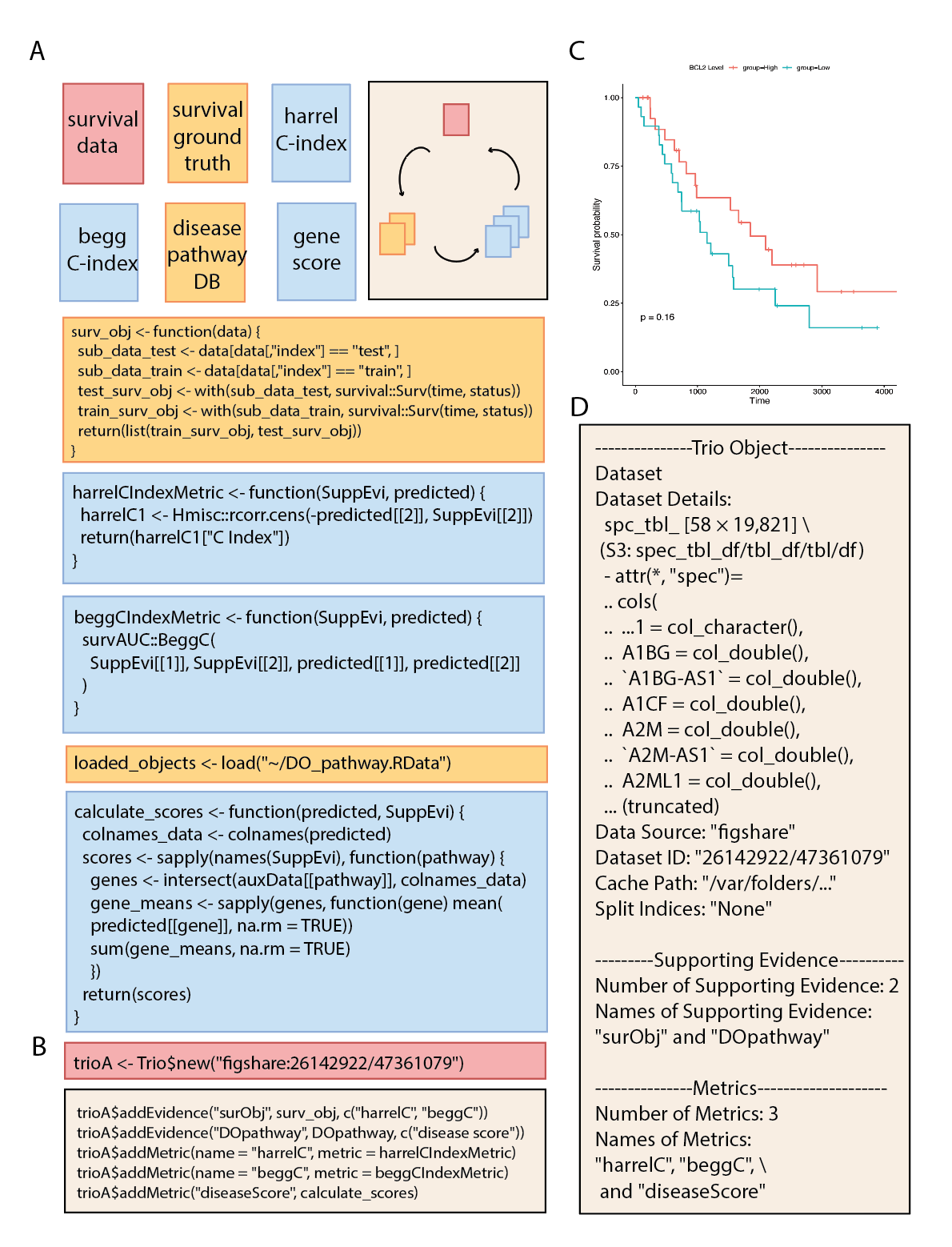

**Supplementary Figure S1. Pseudo code of trio in survival analysis.** A. The color represents different components on the trio object. Red represents the dataset. Yellow represents supporting evidence. Blue represents a metric function. B. The constriction of trio. The trioA is built by the figshare ID. The supporting evidence and metric are added into the trioA. C. Kaplan-Meier curve of ovarian cancer transcriptome data. D. Complete information of trioA object.

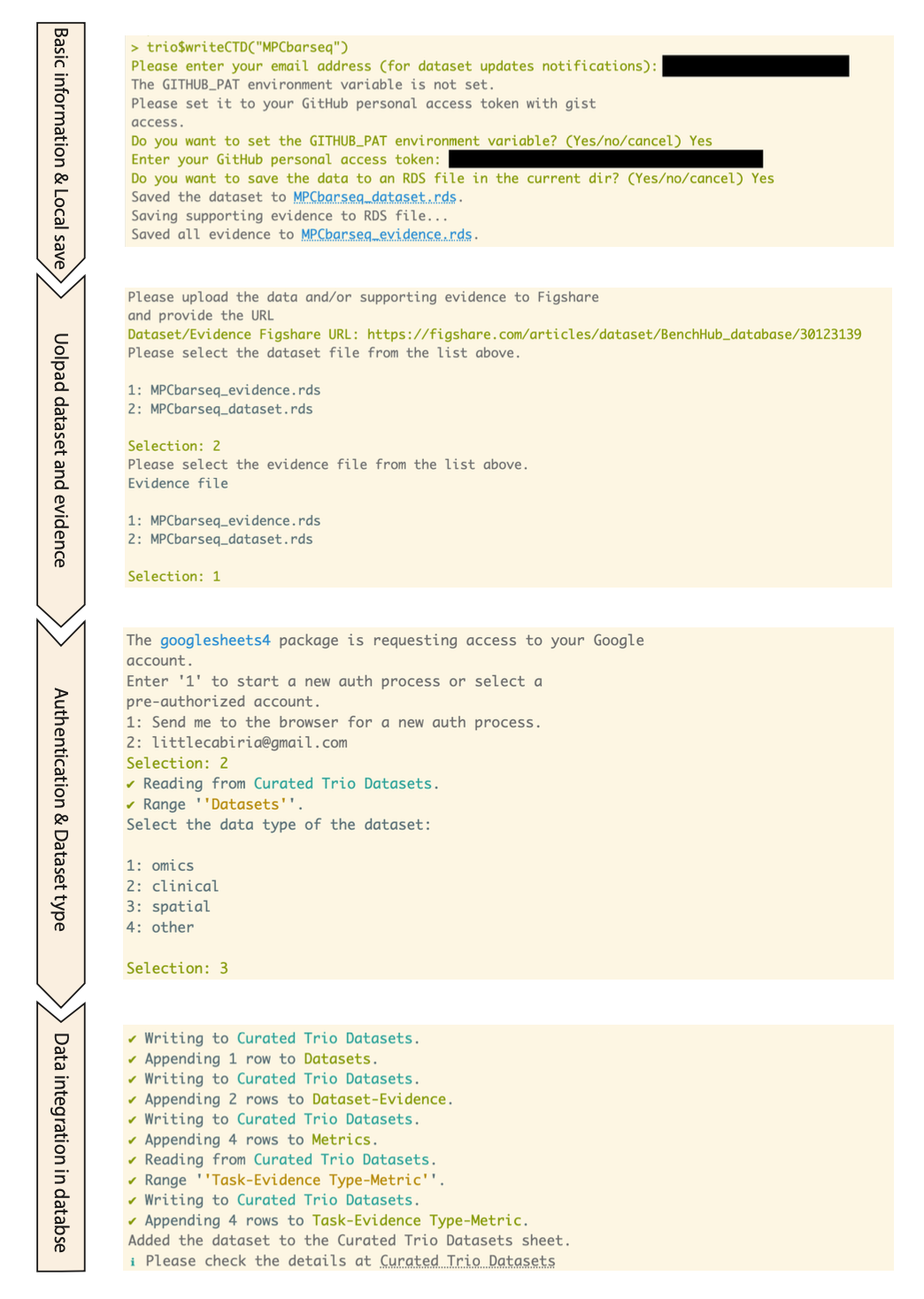

**Supplementary Figure S2. Example workflow for uploading Trio.** The process involves four main steps: (i) provide basic information and save the dataset and evidence files; (ii) upload them to a repository such as Figshare; (iii) authenticating and selecting the dataset type; and (iv) integrating the information into the curated BenchHub Database.
